# Deposited footprints let cells switch between confined, oscillatory, and exploratory migration

**DOI:** 10.1101/2023.09.14.557437

**Authors:** Emiliano Perez Ipiña, Joseph d’Alessandro, Benoît Ladoux, Brian A. Camley

**Author notes:** Please provide details of author contributions here. Please declare any competing interests here.

## Abstract

For eukaryotic cells to heal wounds, respond to immune signals, or metastasize, they must migrate, often by adhering to extracellular matrix. Cells may also deposit extracellular matrix components, leaving behind a footprint that influences their crawling. Recent experiments showed that some epithelial cells on micropatterned adhesive stripes move persistently in regions they have previously crawled over, where footprints have been formed, but barely advance into unexplored regions, creating an oscillatory migration of increasing amplitude. Here, we explore through mathematical modeling how footprint deposition and cell responses to footprint combine to allow cells to develop oscillation and other complex migratory motions. We simulate cell crawling with a phase field model coupled to a biochemical model of cell polarity, assuming local contact with the deposited footprint activates Rac1, a protein that establishes the cell’s front. Depending on footprint deposition rate and response to the footprint, cells on micropatterned lines can display many types of motility, including confined, oscillatory, and persistent motion. On two-dimensional substrates, we predict a transition between cells undergoing circular motion and cells developing an exploratory phenotype. Small quantitative changes in a cell’s interaction with its footprint can completely alter exploration, allowing cells to tightly regulate their motion, leading to different motility phenotypes (confined vs exploratory) in different cells when deposition or sensing is variable from cell to cell. Consistent with our computational predictions, we find in earlier experimental data evidence of cells undergoing both circular and exploratory motion.

**Significance Statement:** Recent experiments showed that epithelial cells modify and sense their local environment, creating a footprint that guides their own motion. Here, we explore how these deposited footprints regulate cell motility. We can recapitulate earlier experimental results with a model that assumes the footprint activates proteins that establish the cell front. We find that cells can use their footprints to change how they explore their surroundings, and that small changes in sensing or depositing footprint can switch the cell from being trapped to being able to explore new environments easily. We find both behaviors in experimental data, suggesting that cells can exhibit multiple crawling behaviors depending on how they deposit and respond to their foot-print.

---

**C**ell migration is a complex process that plays a crucial role in numerous physiological and pathological events, such as tissue development, wound healing, and cancer progression (1). Eukaryotic cells move through the extracellular matrix (ECM) and interact with their surroundings by modulating the ECM through various mechanisms such as deformation, degradation, and deposition of new ECM components (2–4). These actions can produce lasting local changes to the ECM, generating traces like a cellular “footprint.” At the same time, ECM properties such as topography, stiffness, or matrix concentration influence how cells integrate biochemical and mechanical signals from their local environment and modify migration modes, among other physical properties such as shape and speed (5–8). As a result, there is a mutual interplay between cell migration and the ECM facilitated by the traces cells leave behind, allowing cells to sense and follow deposited cues. Eukaryotic examples include neutrophils leaving a trail enriched with the chemokine CXCL12 to direct T-cells to infection sites (9), deposition of vesicular organelles called migrasomes that recruit dorsal forerunner cells that are essential for zebrafish development (10, 11), or migratory fibroblasts guiding cancer cells with a tubular network track on the ECM (12). Bacterial systems also can deposit trails during biofilm formation (13).

Given the coupling between traces cells leave behind and cell motility, this raises a number of questions: Does a single cell’s footprint modify its *own* motion (14)? What characteristics of the cells and their traces control the cell behavior and in particular their migration properties? Can cells change how they interact with the environment to produce different motility behaviors? How can cells sense and respond to footprint signals? Perhaps most importantly: if cells interacting with their footprint is a crucial organizing principle of cell migration, why were signatures of it not seen before the work of (15)?

Previous models addressed the coarse-grained level of how self-propelled objects interact with their trails, including confinement and self-caging in particles that prefer not to cross their own path (15–20). These works focused on the random-walk statistics of self-propelled particles interacting with a deposited footprint, but not how cells sense and respond to the footprint or the role of the imposed pattern geometry and cell shape. We address these questions in this work.

We begin with a focus on recent experiments performed in (15). In this study, the authors found that Madin-Darby canine kidney (MDCK) epithelial cells and Caco-2 human colonic epithelial cells, when placed on fibronectin micropatterned stripes, displayed oscillatory motion resulting from the interaction with their own deposited footprint. Here, using mathematical modeling, we recreate these experiments to examine the motility properties of cells as they interact with the footprint they leave in their path. We can recapitulate the oscillations, but also find other behaviors, including cells that can move persistently if they are able to make their footprint sufficiently quickly. In addition, we use our model to study cell behavior on 2D substrates. One would expect that cells in 2D behave similarly to those on the 1D stripes – if cells are confined by their footprint over stripes, they would also be confined on 2D substrates. Surprisingly, this is not the case. We find that cells that oscillate on 1D stripes exhibit a variety of behaviors on 2D substrates, ranging from being confined and moving in expanding circles to being exploratory with different levels of self-interaction. Such behavior diversity helps explain why interactions with footprints are not as dramatically clear in 2D as in the 1D microstripe environment of (15). We then revisit the experiments of (15) on cells on 2D substrates, and find that cells can develop both expanding circular migration and exploratory migration. Our findings show how the ability of cells to guide themselves using footprints depends on the geometry of the environment in which they move.

## Mathematical model

We build a model to capture the interplay between cell shape, motility, and an extracellular footprint. Our model consists of three interconnected modules (Fig. 1): one for cell shape and motility, one for the dynamics of the footprint, and one for cell polarity – which captures the interaction between the cell motility and the footprint.

**Fig. 1.**
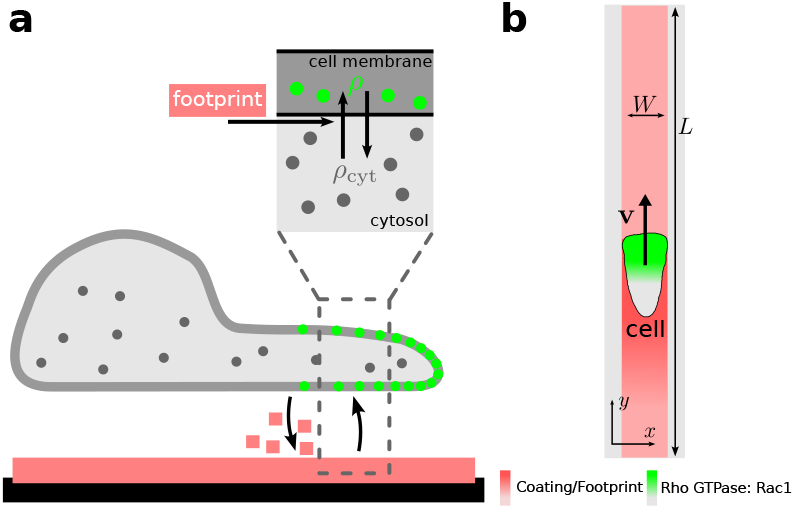
Schematic representation of the model. **a** Side view: cells deposit ECM constituent molecules under their surface that adhere to the substrate, creating a footprint. Cells interact with the footprint which promotes polarization, by upregulating the activation of *ρ*. **b** Top view: linear track of width *W* and large length *L* initially coated with an adhesive substrate *f*_0_ (**r**), such as fibronectin. A polarized cell moves with velocity **v** over the track and leaves a footprint *c*(**r**, *t*) behind.

Cell shape is a crucial factor as it determines both where the cell deposits new footprints and where it can measure signals from the substrate. We use a phase field approach (21– 27) to model the cell’s shape, representing a cell by a field *ϕ*(**r**, *t*) that takes a value of 1 inside the cell and 0 outside. The cell’s boundary is then implicitly defined by *ϕ* = 1*/*2. This approach allows for a simple solution of partial differential equations within the cell’s boundaries (27–30). The cell’s boundary evolves by balancing active forces driving motility and energetic constraints given by a Hamiltonian (22),

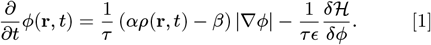

The first term on the right in Eq. (1) is the active propulsion of the cells that acts over the cell membrane. Cell displacement is driven by actin polymerization at the front of the cell, which is defined as the region where a Rho GTPase, which we will assume is Rac1, is in an active state; *ρ*(**r**, *t*) is the concentration of Rac1 in its active state; *ρ*(**r**, *t*) obeys Eq. (5), described later. The cell boundary expands where *αρ > β* and retracts otherwise. The second term on the right of Eq. (1) is the functional derivative of a Hamiltonian representing physical constraints on the cells, such as membrane tension. The Hamiltonian is

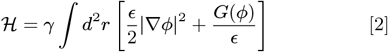

where *G*(*ϕ*) = 18*ϕ*^2^(1−*ϕ*)^2^ is a double well potential with minimums at *ϕ* = 1 (cell interior) and *ϕ* = 0 (cell exterior). *ϵ* sets the thickness of the transition between *ϕ* = 0, 1, and *γ* sets the membrane tension (27).

The second module is the cell’s modification of the local substrate. Earlier work established that these footprints include both cell-produced fibronectin and laminin deposited on the substrate (15), while not ruling out additional components. Depletion of cell-produced fibronectin alone did not stop the oscillations, indicating that the footprint must rely on at least one additional component beyond fibronectin, either laminin or another unidentified matrix component. Following these findings, we consider that cells move over a substrate initially coated with fibronectin *f*_0_(**r**), and that cells deposit an ECM component with concentration *c*(**r**, *t*) onto it, where it remains as a footprint. *f*_0_(**r**) is the initial, potentially space-dependent pattern that the experiments establish by micropatterning (31) and *c* the ECM component – which could be laminin or an as-yet unidentified component – essential to the cell’s response to the footprint. The evolution of the footprint then follows

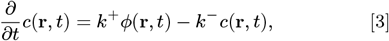

where the cell deposits the ECM component under its surface (represented by *ϕ*) with deposition rate *k*^+^ and degrades with rate *k*^*−*^. The concentration of *c* is measured in terms of a reference concentration unit *c*_0_ – deposition rate *k*^+^ has units of *c*_0_*/*s.

The last ingredient in our model is how cells interact with the footprint. Cells adhere to the ECM through integrins, transmembrane proteins that have mechanosensitive roles mediating inside-out and outside-in signaling pathways (1, 32– 35). Thus, we assume that the cell senses and responds to the footprint via an integrin-mediated pathway, represented by an effective response function *ψ*(**r**, *t*), which will later regulate the cell’s biochemical polarity, changing *ρ*(**r**, *t*). The response function *ψ* takes values between 0 to 1, with 0 being where there is no footprint and 1 where there is enough footprint to saturate the response. We write the effective response function *ψ*(**r**, *t*) as a Hill-like equation,

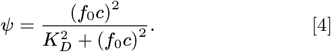

Eq. (4) assumes that to have a response to the footprint (*ψ >* 0) requires the presence of *both* the adhesive compound *f*_0_ and the deposited footprint *c*(**r**, *t*).

The cell’s biochemical polarization – what directions it chooses to protrude, and how it responds to the presence of the footprint – is modeled by an extension of the simple Rho GTPase model of (36), in which there is a membrane-bound active form of Rac1 that promotes actin polymerization, *ρ*(**r**, *t*), and a non-active cytosolic form *ρ*_cyt_(*t*). Rac1 can diffuse on the membrane and in the cytosol, and can exchange between the two states, following a reaction-diffusion scheme (Fig. 1**a** box). We note that our model of the cell, following e.g. (26, 30) and others, represents a 2D section of the cell membrane, which can be thought of as the basal surface or a 2D projection of the 3D cell; hence even the active Rac1 is localized over the entire region *ϕ >* 0. The active Rac1 concentration *ρ* then obeys

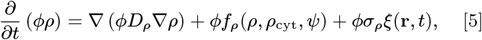

where *D*_*ρ*_ is the membrane diffusion coefficient of the active Rac1, *f*_*ρ*_ is a reaction term describing Rac1 activation and inactivation, *σ*_*ρ*_ parameterizes the intrinsic fluctuations in Rac1 activity and *ξ*(**r**, *t*) is an uncorrelated white noise with ⟨*ξ*(**r**, *t*)*ξ*(**r**^*′*^, *t*^*′*^)⟩ = *δ*(**r**−**r**^*′*^)*δ*(*t*−*t*^*′*^) and *δ* the Dirac delta function. The *ϕ* field in Eq. (5) acts as a function indicating the cell interior; this approach models *ρ* as diffusing only within the cell (*ϕ >* 0), simplifying the handling of complex boundary conditions (28, 29).

For the reaction term *f*_*ρ*_, we use a modified version of the model of (36),

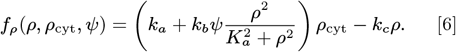

where *k*_*a*_ is the basal activation rate of *ρ, k*_*c*_ the basal de-activation rate, and the term proportional to *k*_*b*_ is a positive feedback mechanism that promotes Rac1 activation where *ρ* is large; *K*_*a*_ sets the level of *ρ* where this positive feedback begins to saturate. This positive feedback term leads to a solution of Eq. (5) where *ρ* is stably polarized across the cell, with high *ρ* setting the front of the cell and low *ρ* in the back (36). We extend the model of ref (36) by assuming the positive feedback term is proportional to the effective footprint *ψ*. This is our core assumption about how the cell senses the footprint. Our motivation for this is based on the observation that when integrins bind to the matrix, they activate pathways that promote actin polymerization by mediating Rac1 signaling, thereby facilitating directed cell migration (32) in a way akin to haptotaxis (1). Thus, when the cell moves over a previously visited region where it has already laid out a footprint, the positive feedback mechanism becomes active, leading to a local increase in *ρ*(**r**, *t*). On the other hand, if the cell reaches an unexplored region where no footprint is present, the positive feedback mechanism is deactivated, causing *ρ*(**r**, *t*) to decrease. This results in the cell being more likely to move away from the unexplored areas.

The non-active cytosolic Rac1, *ρ*_cyt_, diffuses faster than the membrane-bound active one, and we treat it as well-mixed (spatially uniform),

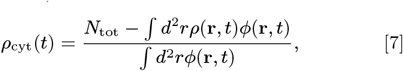

where *N*_tot_ is the total number of Rac1 proteins inside the cell.

## Results

### Motility Patterns and the Mechanism of Oscillations

To mimic the experiments of (15), we set the fibronectin concentration *f*_0_(**r**) to be uniformly coated linear tracks of width *W* = 10*µ*m and large length *L* = 500*µ*m, Fig. 1**b**. We placed a circular cell of *R* = 10*µ*m in the center of the track over a small rectangle of height 2*R* and width *W* coated with *c* = 100 *c*_0_ and let it settle for 50 s while relaxing to its preferred shape. The initial rectangle provides a suitable substrate so the cell can take a natural shape before starting the simulation. During this setup time, the cell does not deposit any footprint, and Rac1 concentration *ρ* is kept uniform. Then we induce the cell polarization in a random direction, turn on Rac1 and footprint dynamics, given by Eq. (5) and Eq. (3) respectively, and let the system evolve for 60h or up to when the cell reaches the edges of the track. Cells can display a variety of motility patterns, depending on their parameters (Table 2), including cells moving persistently in one direction (Mov. 1), cells oscillating with increasing amplitude (Mov. 2), cells diffusing while they oscillate (Mov. 3) to cells that are unable to polarize and, thus do not move (Mov. 4), see Fig. 2**a**. Three of these behaviors – unpolarized cells, cells with increasing oscillations, and persistent cells – were observed in the experiments of (15) in different circumstances.

**Fig. 2.**
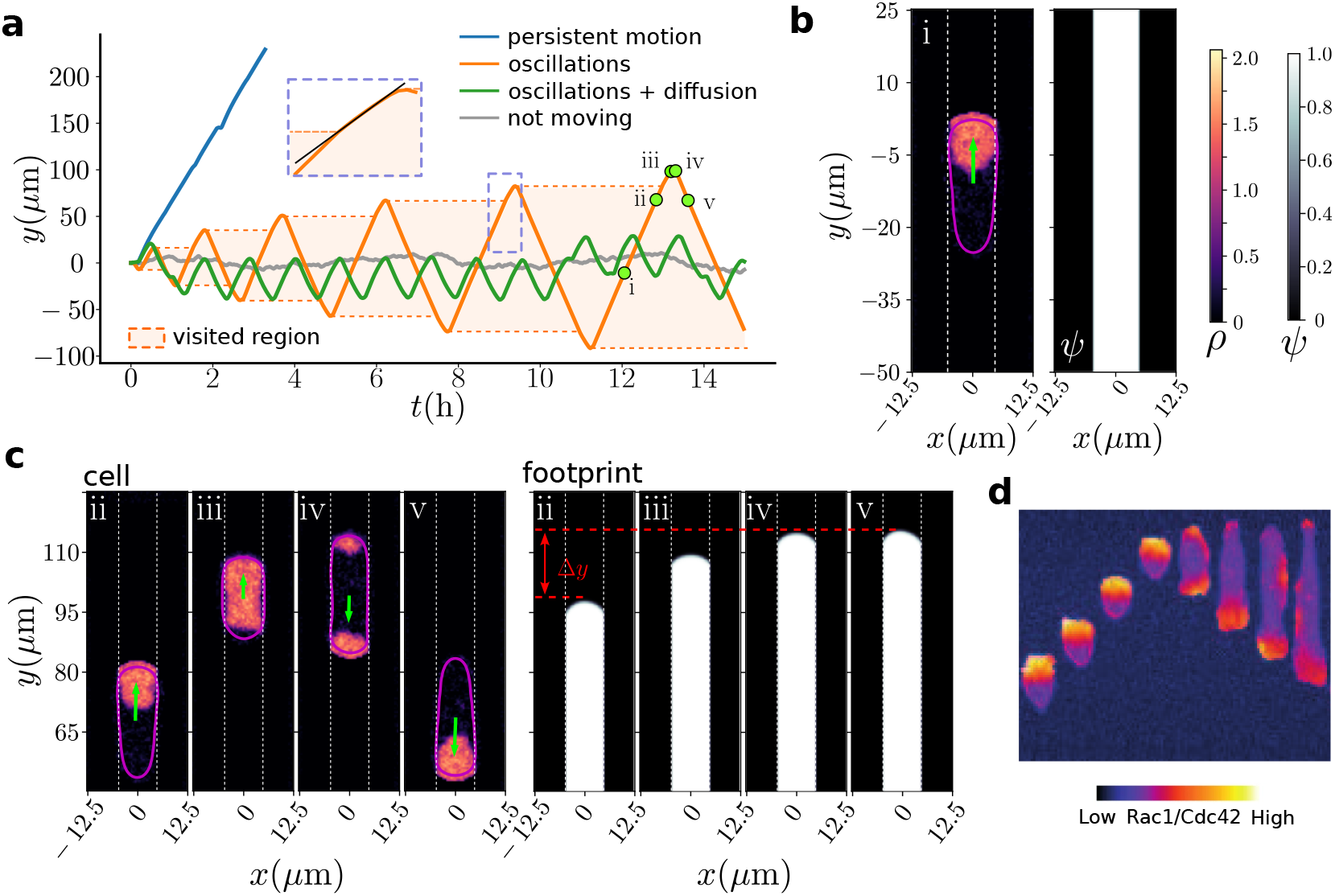
**a** Example of trajectories corresponding to different motility behaviors of the model: depending on the parameters we observe persistent motion (*k*^+^ = 0.6 *c*_0_ */*s), oscillations with increasing amplitudes (*k*^+^ = 0.3 *c*_0_ */*s), cells diffusing as they oscillate (*k*^*−*^ = 7.2 h^*−*1^), and cells stalling without being able to move (*σ*_*ρ*_ = 0.47 *µ*m^*−*2^ s^*−*1*/*2^, *k*_*b*_ = 8 s^*−*1^). Parameters not specified are the defaults listed in Table. 2. The light orange shaded area represents the regions the cell visited in the case of the oscillation with increasing amplitudes. In the inset, a trajectory is amplified to illustrate a slight change in slope and the slow down of the cell velocity when it reaches the edge of the visited region. **b** Snapshot of a simulation corresponding to the point i in panel **a**. Left: a cell in motion. The cell border is marked in purple. Rac1 (*ρ*) is high at the front of the cell. Right: effective footprint, *ψ*. **c** Snapshots showing a cell reaching the edge of the footprint (points ii, iii, iv, and v in panel **a**). Left: the cell slows down and repolarizes reversing the direction of motion. Right: during this process, the footprint extended a distance Δ*y*. The region enclosed by the dashed white lines in panels **b** and **c** represents the tracks initially coated with fibronectin. **d** Kymograph of an MDCK cell expressing the Rac1/Cdc42 activation reporter PBD-YFP during a reversal event that occurs when the cell reaches the edge of the explored region. The image is cropped from Fig. 1i of (15).

We initially focus on the oscillation with increasing amplitude patterns (Fig. 2**a**, orange curve). We observe that cells move with constant velocity when they are on a past trace (light orange shaded area in Fig. 2**a**), and slow down (change in the slope of the *y*(*t*) curve, see inset in Fig. 2**a**) when they enter unexplored regions, then finally stop and reverse the direction of motion. To further understand this, we mark 5 points in the orange line in Fig. 2**a** representing different moments on the trajectory of the cell. The first one, (i), shows the cell moving in the center of the footprint. At this position, the cell is polarized, showing an accumulation of *ρ* at the front (Fig. 2**b** left panel). The cell has crossed this region repeatedly, saturating the substrate with its footprint, so *ψ*≈ 1, as shown in the right panel in Fig. 2**b**. The other 4 points, (ii)-(v), show the cell in the process of reaching the edge of its footprint and turning around. At points (i) and (ii), the effective footprint is saturated (*ψ*≈1) both behind and in front of the cell – the cell feels no local haptotactic gradient tending to repolarize it. At point (iii), the cell’s front enters into a previously unexplored region. *ρ* then decreases at the cell front, leading to the front’s velocity decreasing; the cell then decreases in size as the back catches up. This decrease in velocity arises because in the unexplored region, where *ψ*≈0, the positive feedback mechanism that sustains polarity at the front no longer holds, leading to decreased *ρ* via Eq. (6). At point (iv) we see that there are two competing protrusions (regions of high *ρ*) – the original protrusion, and a new one pointing toward the region of high footprint – here, the direction from which it came. These protrusions temporarily compete, but eventually the protrusion at higher *ψ* wins, and the cell turns around. Finally, at point (v) we see the cell traveling on its previous trace over a fully developed footprint. In the right panel of Fig. 2**c** we can see that during cell flipping, the footprint was extended a length of Δ*y*. This indicates that the next time the cell arrives at this edge it will reach a further distance Δ*y* than before, resulting in oscillations with increasing amplitudes. Moreover, during each cycle, the cell extends the footprint another Δ*y*, showing that the amplitudes should eventually, after an initial transient, increase linearly with time, see Fig. 3**a**. (The linear increase in amplitude with time is true only as long as the footprint degradation rate is zero, *k*^*−*^ = 0, as we will discuss later.)

**Fig. 3.**
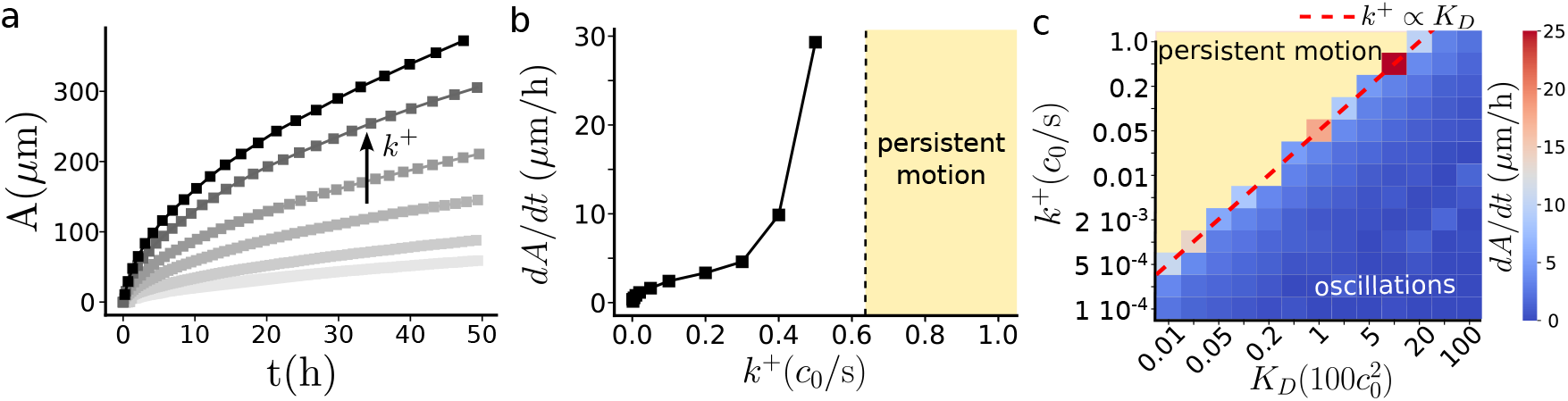
Footprint deposition dynamics controls motility behavior. **a** Oscillation amplitude *A* increases with time and with the footprint deposition rate *k*^+^. **b** Rate of change of amplitude over time as a function of the footprint deposition rate. Beyond a characteristic deposition rate cells transition from oscillatory to persistent motion. **c** Phase diagram of amplitude rate of change as a function deposition rate and *k*_*D*_. The transition from oscillatory to persistent motion (yellow area) is marked in a red dashed line and follows *k*^+^∝*K*_*D*_. The rate of change of *A* was calculated as the slope of a linear fit over the last five periods of the curves in panel **a**. All simulations were done using default parameters in Table 2 with the exemption of the deposition rate, *k*^+^, and the effective footprint response threshold *K*_*D*_ in panel **c**. In panel **a** the curves corresponds to *k*^+^ = {0.01, 0.02, 0.05, 0.1, 0.2, 0.3} *c*_0_ s^*−*1^.

Measurements of Rac1 activity by d’Alessandro et al. (15) on cells reversing at the end of their oscillations also see repolarization of Rac1 as the cell reaches the end of the footprint. These measurements, in agreement with our theory, show a transient state where both the front and back of the cell have elevated Rac1 activity; these data are reproduced in Fig. 2**d**.

One key finding in the original work of (15) was that footprint increased persistent motility. This was discovered using “conditioned” substrates, where cells were allowed to crawl over substrates at high density for a long time, leading to the substrate being saturated with footprint. After removal of the original cells, when new cells were placed on the conditioned substrate, rather than oscillating as before, they moved in a highly persistent manner. We also observe in our simulations that cells move highly persistently when they crawl over a substrate that has been saturated with the footprint, if we set *ψ* ≈ 1 everywhere, Mov. 5.

### Footprint deposition dynamics controls existence and amplitude of oscillations

How does motility depend on the rate at which the cells deposit and generate the footprint, which is necessary for cells to crawl and move? We explore cell motion for different footprint deposition rates, *k*^+^. We initially assume the footprint does not degrade with time, *k*^*−*^ = 0, supported by the experiments of (15) showing that conditioned substrates still induce persistent motion after *>* 24 hours. Increasing the deposition rate, we find that cells oscillate with larger amplitudes *A*, (Fig. 3**a**). Depositing more footprint allows cells to form the necessary substrate faster. The effect is not noticeable when the cell crawls over past positions, as the footprint gets rapidly fully developed and saturated (*ψ*≈1). Thus, the cells move over their previous traces with the same velocity almost independently of their footprint deposition rate *k*^+^. But as cells reach the edge of the footprint, their ability to build more substrate faster allows them to travel farther into unexplored regions before stopping and turning around.

Surprisingly, as we increase the deposition rate *k*^+^ further, we do not simply see a continued increase in oscillation amplitude, but we see a sudden transition to a persistent motion, where cells no longer oscillate. We visualize this by plotting *dA/dt* as a function of deposition rate *k*^+^ (Fig. 3**b**). Persistent motion occurs when cells can deposit enough material to maintain polarization before the haptotactic cue reverses their polarization. For our default parameters, the transition to persistent motion occurs at *k*^+^ ∼0.6*c*_0_*/s*, but the transition value also depends on *K*_*D*_, which controls how much footprint is necessary for the cell to reach its full response. Varying *K*_*D*_ and *k*^+^ simultaneously, we see that higher values of the threshold *K*_*D*_ promote the oscillatory behavior of the cells, requiring higher footprint deposition rates for the transition into persistent motion (Fig. 3**c**), as we would expect. The dashed transition line on the phase diagram in Fig. 3**c** has the form *k*^+^∝*K*_*D*_. This relationship is predicted by a simple model we introduce later (Eq. 9).

Past experiments have also observed transitions between persistent motion and oscillations given various biochemical interventions on cells, including zyxin knockout and microtubule destabilization (37–39). In these cases, the underlying mechanism driving the oscillations was not believed to be induced by interaction with the substrate, but by perturbation of the cell’s internal machinery or competition for myosin (39). However, our results suggest that changes in deposition, degradation, or haptotactic sensing could also mediate similar transitions.

### Footprint aging confines cells and sets a memory length

In (15), it was reported that cells leave a long-lived footprint, suggesting that the substrate does not degrade significantly during the experimental times ∼60*h*. Nevertheless, *in vivo* we expect ECM components to be actively degraded by matrix metalloproteinase enzymes (MMPs), whose misregulation is associated with various diseases, including tumor progression (40). How does footprint aging alter single-cell motility? If the footprint degrades, a cell will lose the memory of its previous trajectory – if it moves too far away from its old path, the path will no longer be present. Cells performing the expanding oscillation of Fig. 2**a** thus cannot increase their oscillation amplitude indefinitely – eventually they will not reach their past footprint before it decays. Simulations with footprint degradation show cells reach a steady-state oscillation amplitude (Movie 3). We performed simulations for varying values of footprint degradation rate *k*^*−*^ (Fig. 4**a**), choosing parameters leading to cells that oscillate with increasing amplitude when *k*^*−*^ = 0. When the footprint degradation rate is non-zero, cells oscillate with increasing amplitude until the amplitude reachs a steady-state value *A*_ss_ (Fig.4**a**).

**Fig. 4.**
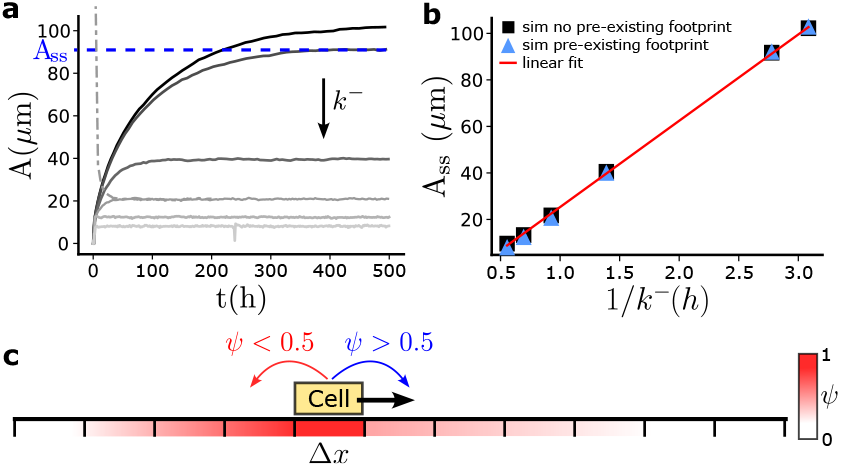
Footprint aging causes cell confinement. **a** Amplitude as a function of time for different footprint degradation rates. In case of footprint degradation, cells adjust their amplitude until reaching a steady-state oscillation with amplitude *A*_ss_. Solid lines correspond to cells starting from an initial condition without a preexisting footprint. The dashed-dotted line is an example of a cell starting from a preexisting footprint larger than the steady-state amplitude value, *A*_ss_. **b** Steady-state amplitudes as a function of the inverse of the degradation rate. Simulations in black squares have no preexisting footprint and simulations in blue triangles are initialized with a preexisting footprint. The red line is a linear fit. **c** Schematic representation of the simplified model of a cell moving over a 1D lattice. A cell continues to move in the same direction if *ψ >* 0.5 and changes the direction of motion otherwise. Parameters: *k*^+^ = 0.01 *c*_0_ s^*−*1^ and *k*^*−*^ = {0.324, 0.36, 0.72, 1.08, 1.44, 1.8} h^*−*1^.

The faster the substrate and footprint degrade, the smaller the steady-state amplitudes are and the faster this steady-state condition is reached, see Fig. 4**a**. Higher degradation rates means the cell has less time to revisit a place before the memory of previous arrivals is lost, *i*.*e*. shorter memory length and greater confinement. We quantify this in Fig. 4**b** where we show the steady-state amplitude values *A*_ss_ as a function of the degradation characteristic time, 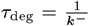.

Interestingly, the steady-state amplitudes are proportional to the degradation time, suggesting that the value of *A*_ss_ arises from a cell moving at a constant speed for a time proportional to the degradation time.

The steady amplitude is robust to changes in the initial state of the substrate. In simulations where cells are initiated on a preexisting footprint (coated on 2/3 of the box length with *c* = 100 *c*_0_ so that *ψ*≈1), cells oscillate with decreasing amplitudes as the footprint decays until they reach a steady-state (dashed-dotted line in Fig. 4**a**). This final oscillation has the same amplitude as in the case of the linear track without the preexisting footprint (blue triangles in Fig. 4**b**).

To gain more insight into what is controlling the different cell motility behaviors and in particular the oscillations, we develop a simplified model. Our previous simulations and the experiments of (15) suggest that cells move with roughly constant speed within an existing footprint. We thus build a simple model of a cell moving at constant speed on a 1D lattice, see Fig. 4**c**. In this simplified 1D model, the deposited footprint is composed of a single component that is deposited by the cell and degrades with time. The cell always moves with constant speed 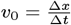 and follows simple rules: after each time step Δ*t* if the local footprint concentration under the cell position is below a threshold value 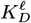, then the cell changes direction and returns to the previous lattice site; otherwise, it moves to the next lattice site in the direction it was originally traveling. We use a superscript/subscript *l* to indicate “lattice” variables, which will have different units than their corresponding values in the full simulation. The time step Δ*t* reflects the time the cell takes to make a decision – this would depend on, e.g. the detailed parameters of the Rac1 reactions within our more complicated model. A cell will move in one direction until reaching a position where the footprint is low enough to make the cell turn around and start moving in the opposite direction. We write the footprint concentration dynamics for the simple model as

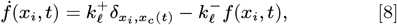

where 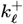 and 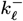 are the deposition and degradation rates, 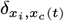 is the Kronecker delta function and *x*_*c*_(*t*) is the position of the cell at time *t*. To compute the steady-state amplitude *A*_ss_, we consider a cell that has already reached the stationary state. To maintain a steady state, the footprint concentration at the boundaries must remain the same during a full loop of the cell, *i*.*e*. the time between when the cell departs and returns to the edge. Since the boundary is where the cells turn around, then the footprint concentration at that position must be equal to 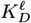. During an oscillation period, the total amount of deposited footprint at the boundary is 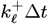. The time required for a cell to complete an oscillation (travel a distance 2*A*_ss_) is just 2*A*_ss_*/v*_0_, so the amount of footprint lost at the edge during a complete cycle is 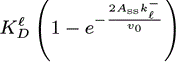. At steady state, the amount of footprint added at the edge must equal the amount degraded, so we can equate 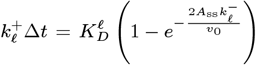. We then find the steady-state amplitude as

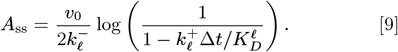

This expression sheds light on the role of the different model parameters. For instance, Eq. (9) shows analytically that steady-amplitude is linearly proportional to degradation time, 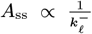, as observed in the more complex simulation Fig. 4**b**. Similarly, Eq. (9) reveals that faster cells have larger amplitudes since they can cover larger distances before the footprint degrades below the threshold concentration. The most remarkable prediction from Eq. (9) is the transition from oscillation to persistent motion. The logarithm in Eq. (9) diverges when 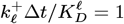, indicating the moment when cells become persistent. This predicts that the transition line should have *K*_*D* ∼_*k*^+^ – as observed in our complex simulation (Fig. 3**c**). Essentially, we expect that cells will be persistent, roughly speaking, when they deposit enough footprint in time Δ*t* to reach the threshold of footprint response 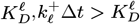. This condition for persistent migration is still reasonable in the absence of degradation (*k*^*−*^ = 0), as in our simulations in Fig. 3.

This simple model is remarkably effective at capturing the behavior of cells on 1D micropatterned stripes, predicting correctly the transition between oscillatory and persistent motion seen in Fig. 3**c**, as well as the dependence of amplitude on footprint decay rate (Fig. 4**b**). However, it naturally cannot capture the full complexities of cell and footprint geometry. This complex geometry becomes primarily relevant when we simulate cells on two-dimensional substrates with uniform fibronectin, *i*.*e*. unconfined cells.

### Cells in 2D environments transition between circularly– confined and exploratory motion

Cells in 1D tracks are constrained to only move in two directions. How do cells behave when they have more freedom to move? Here, we chose the initial fibronectin micropattern *f*_0_(**r**) to be 2D squares of edge length *S* = 250*µ*m and then placed a cell in the center. We explored cell trajectories for varying values of the footprint deposition rate *k*^+^ (Fig. 5). As in the first part of this work, we assume no degradation, *k*^*−*^ = 0. At small deposition rates, cells are confined and need to build their substrate to move around (small *k*^+^ in Fig. 5**ab**). This is similar to our observations on linear stripes for small deposition rates, though in the 2D confinement we do not observe oscillations with increasing amplitudes but instead cells moving in circles slowly expanding their explored area, see Fig. 5**a** and Movie 6. Increasing *k*^+^ further, we see that cells cover more explored area in less time (compare top left panel and right panel in Fig. 5**a**).

**Fig. 5.**
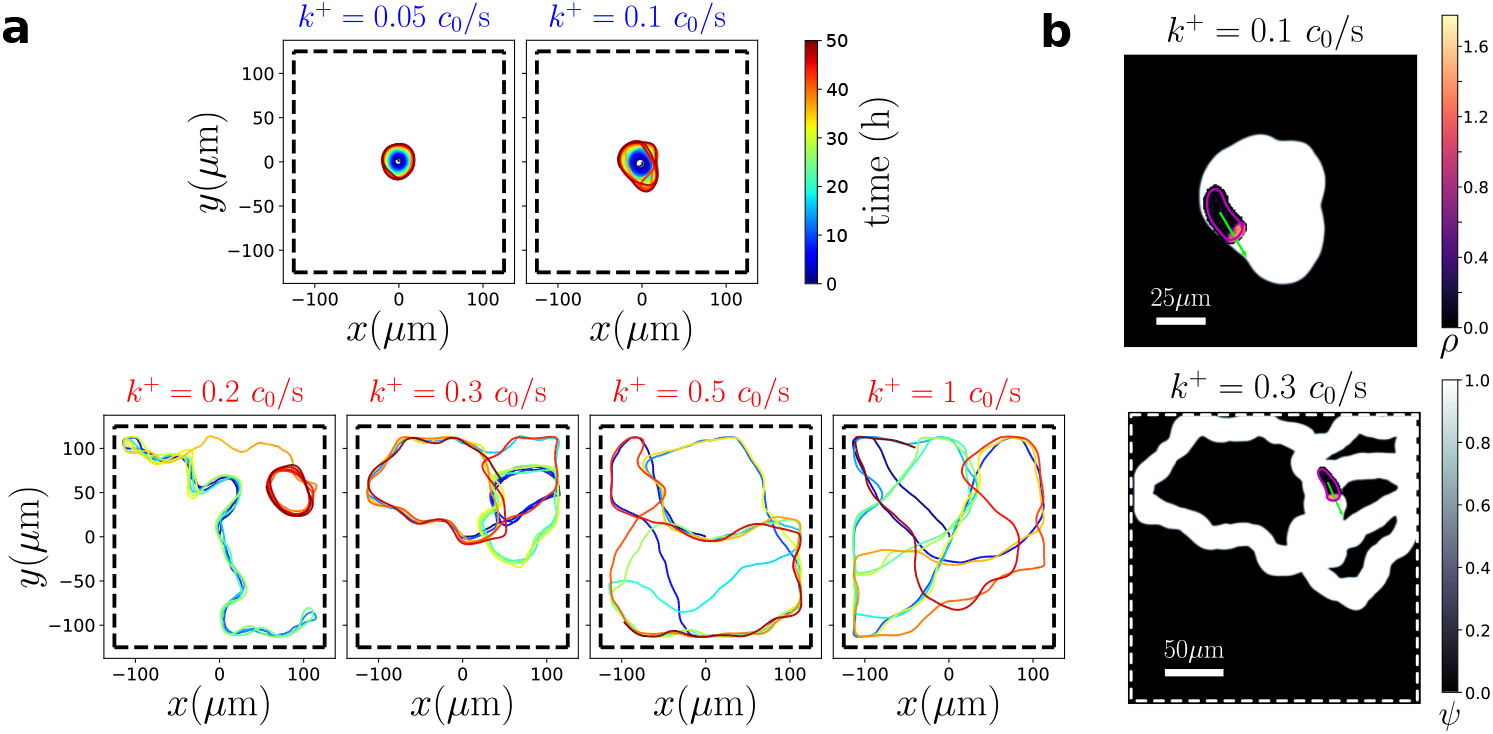
Cells over 2D substrates show transitions between highly confined motility to a complex self-interacting exploratory behavior. **a**. Cell trajectory examples for different deposition rates. Color code indicates time. Top (bottom): examples for a deposition rate below (above) the motility behavior transition. **b**. Snapshots of movies showing the shape, polarity (*ρ*), and velocity of the cell (green vector), along with its footprint traces (*ψ*) for a cell secreting below (top – Mov. 6) and above (bottom – Mov. 7) the transition. The dashed lines indicate the boundary of the fibronectin-coated region (*f*_0_ (**r**))

If we keep increasing the deposition rate *k*^+^ further we again, similarly to the 1D case, observe that there is a transition from a highly confined motility behavior to a more persistent and exploratory one (Fig. 5). For these large deposition rates (Fig. 5, bottom), we observe cells at first moving like persistent random walkers, Mov. 7. When cells randomly exploring the free space run into their previous trajectory, they tend to reorient their direction toward the previously-explored space and start moving on the footprint they have created, see Fig. 5**a** lower panels. This produces a sort of weak confinement, in which cells can in principle freely explore all available space, but are most likely to be found in previously visited regions. Cells are more likely to remain on the prior trajectory for smaller values of the deposition rates, *k*^+^. Conversely, as cells deposit more footprint they can more easily escape from previous trajectories and become less confined. Table 1 summarizes the different migration modes in 1D and 2D regarding their confined/exploratory features.

**Table 1.** Summary of migration modes in 1D (striped coating) and 2D (square coating).

|  | 1D | 2D |
| --- | --- | --- |
| (loosely) confined | **Oscillatory motion**<br>cells oscillate (with increasing amplitude) within the previously visited areas. | **Circular motion**<br>cells move in (expanding) circles around the edge of the previously visited areas. |
| exploratory | **Persistent motion**<br>cells move persistently in one direction being able to migrate into unvisited regions. | **Self-interacting exploratory behavior**<br>cells alternate between exploring new areas and moving along their previous trails. |

**Table 2.** Parameter selection. Most of the parameter values were extracted from (22) and then changed by rescaling time to ensure the velocity of the cell matched the experiments. Unless otherwise indicated in the text, these are the parameters used throughout the results.

| Parameter | Description | Value |
| --- | --- | --- |
| $\Delta r$ | Space grid | $0.5\mu\text{m}$ |
| $\Delta t$ | 1D simulation time step | $0.05\text{s}$ |
| | 2D simulation time step | $0.2\text{s}$ |
| $\Delta t_{\text{data}}$ | 1D data storage time | $250\text{s}$ |
| | 2D data storage time | $400\text{s}$ |
| $R_{\text{cell}}$ | Initial cell radius | $10\mu\text{m}$ |
| $W$ | Linear track width | $10\mu\text{m}$ |
| $L$ | Linear track length | $500\mu\text{m}$ |
| $S$ | 2D square substrate length Fig. 5 | $250\mu\text{m}$ |
| | 2D square substrate length Fig. 6-7 | $1000\mu\text{m}$ |
| Footprint |  |  |
| $c_0$ | Arbitrary concentration units | — |
| $f_0$ | Adherent coating | $1000 c_0$ |
| $K_D$ | Effective footprint dissociation constant | $1000 c_0^2$ |
| $n_{\text{hill}}$ | Hill coefficient | 2 |
| $k^+$ | Footprint deposition rate | $0.3 c_0/\text{s}$ |
| $k^-$ | Footprint degradation rate | $0 \text{s}^{-1}$ |
| Phase field |  |  |
| $\alpha$ | Protrusion strength | $0.8 \text{ pN } \mu\text{m}$ |
| $\beta$ | Retraction strength | $0.1 \text{ pN } / \mu\text{m}$ |
| $\gamma$ | Cell tension coefficient | $1.8 \text{ pN}$ |
| $\epsilon$ | Interface width | $2 \mu\text{m}$ |
| $\tau$ | Friction coefficient | $26.2 \text{ pN s } / \mu\text{m}^2$ |
| Cell's polarity |  |  |
| $N_{\text{Tot}}$ | Total amount of Rac1 | 600 |
| $D_\rho$ | Membrane $\rho$ diffusion constant | $0.08 \mu\text{m}^2/\text{s}$ |
| $\sigma_\rho$ | Fluctuation intensity in $\rho$ concentration | $0.22 \mu\text{m}^{-2} \text{s}^{-1/2}$ |
| $k_a$ | Rac1 activation rate | $0.01 \text{s}^{-1}$ |
| $k_b$ | Rac1 positive feedback activation rate | $1 \text{s}^{-1}$ |
| $K_a$ | Rac1 positive feedback threshold | $1 \mu\text{m}^{-2}$ |
| $k_c$ | Rac1 inactivation rate | $1 \text{s}^{-1}$ |

The experimental measurements of (15) for cells on microstripes are in the confined limit – they show oscillations of increasing amplitude. Would we also expect cells on a uniformly-coated 2D substrate to be confined? Our simulations in Fig. 6 show the answer to be: not necessarily! The transition between confined motion and an exploratory one occurs in a different range of values of deposition rates in 1D and 2D. For our default parameters the transition in 1D occurs in the range 0.6 *c*_0_*/*s *< k*^+^ *<* 0.7 *c*_0_*/*s while in 2D, it is 0.1 *c*_0_*/*s *< k*^+^ *<* 0.2 *c*_0_*/*s (Fig. 6, Fig. 3**b**, Fig. 5**a**)). This means it is possible to have the same cell with the same parameters have a confined oscillatory motion in 1D but undergo exploration in 2D (Fig. 6). We should also note that 2D migration is more sensitive to changes in deposition rate *k*^+^ than 1D migration – there is a wide range of deposition rates where cells have confined oscillations in 1D, while exhibiting significant variability in their self-interacting exploratory levels in 2D (points *k*^+^ = 0.1, 0.2, 0.3*c*_0_*/s* in Fig. 6). This suggests that if there is cell-to-cell variability in deposition rate, it might be more apparent on 2D substrates, while the confined oscillation may be more universal.

**Fig. 6.**
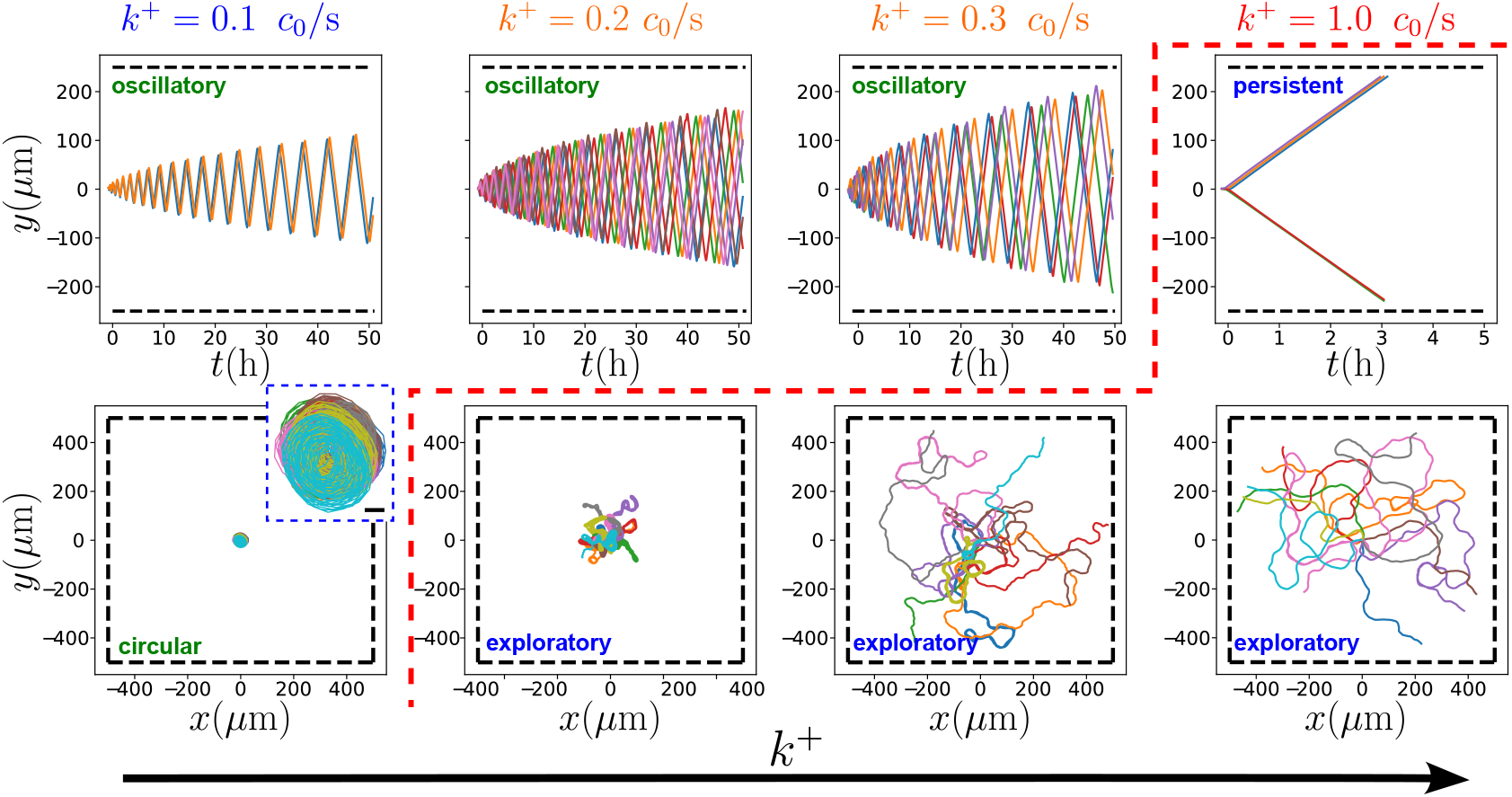
Transitions between confined and persistent cells occur at different values of the deposition rate on 1D striped and 2D squared substrates. Each panel shows a set of 10 trajectories arranged by increasing values of *k*^+^ from left to right. Top panels correspond to stripes and bottom panels to squares. In the lower-left corner, the inset shows a magnification of the trajectories with a scale bar of 10*µ*m. The dashed red line marks the transition between confined and persistent movement. In this figure, we increased the polarity noise to *σ*_*ρ*_ = 0.32 *µ*m^*−*2^ s^*−*1*/*2^ to more closely match the roughness of the simulated trajectories to the experimental ones. In the bottom panels, cells are simulated on a 2D square substrate of edge length *S* = 1000 *µ*m and the simulation stopped when the cells reach the edge.

Why is the transition point from confined to exploratory motion so different in 2D than in 1D in Fig. 6? This quantitative change may reflect the cell’s shape. Cells in 2D moving over unexplored substrates tend to be more rounded than cells on the 1D stripes. In the language of our simplified model, cell shape may alter the effective deposition rate for a cell 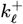 (since the cell’s area changes), or the time to make a decision Δ*t*. To verify the effect of cell shape, we performed simulations of cells with different tension coefficients *γ*, and found that changing *γ* can change cells from confined to exploratory (Fig. S1). However, the effect of *γ* is not monotonic – increases and decreases in *γ* can both create this transition. This is similar to non-monotonic effects of cell tension on reorientation we previously discovered in (30). We also explored the possibility that the early transition in 2D is driven by the polarity noise, *σ*_*ρ*_. However, we found that the transition point is robust to an increase in noise, on both, 1D stripes and the 2D substrates, see Fig. S2.

### Experiments observe mixture of circular confined motion and exploratory motion

Our model predicts (Fig. 6) that cells which oscillate on 1D stripes – like the MDCK cells of (15) – can either develop circular trajectories or exploratory ones on 2D substrates, depending on the deposition rate. What happens experimentally? We show selected experimental trajectories from (15) of MDCK cells on a 2D fibronectin substrate in Fig. 7**a** and the full set (*n* = 200) in Fig. S3.We do see some cells moving in circles with an increasing area, consistent with cells being below the transition point as in the case of the 1D stripes, Fig. 7**a** top panel. However, these were rare cases. We classified trajectories as circular or exploratory, using an initial algorithmic filter and then a blinded categorization (Section S3 in the SI). Only 12 out of 150 classified cells showed circular trajectories, and, in most cases, they eventually escaped from the confined circular motion to start exploring new areas. The vast majority of cells showed exploratory behavior (138). These trajectories show occasional backtracking and crawling over a pre-existing path, see Fig. 7**a** bottom. An additional 50 cells had trajectories ambiguous enough that we did not classify them. This contrasts with the absence of cells with circular behavior found on 154 classified trajectories on conditioned substrates (Fig. S4) – suggesting these circular trajectories are consequences of the footprint deposition. Together, these data suggest that real MDCK cells are close enough to the transition point between confined and exploratory motion in 2D that cell-cell variability in deposition rates or cell shape allows them to undergo both behaviors.

**Fig. 7.**
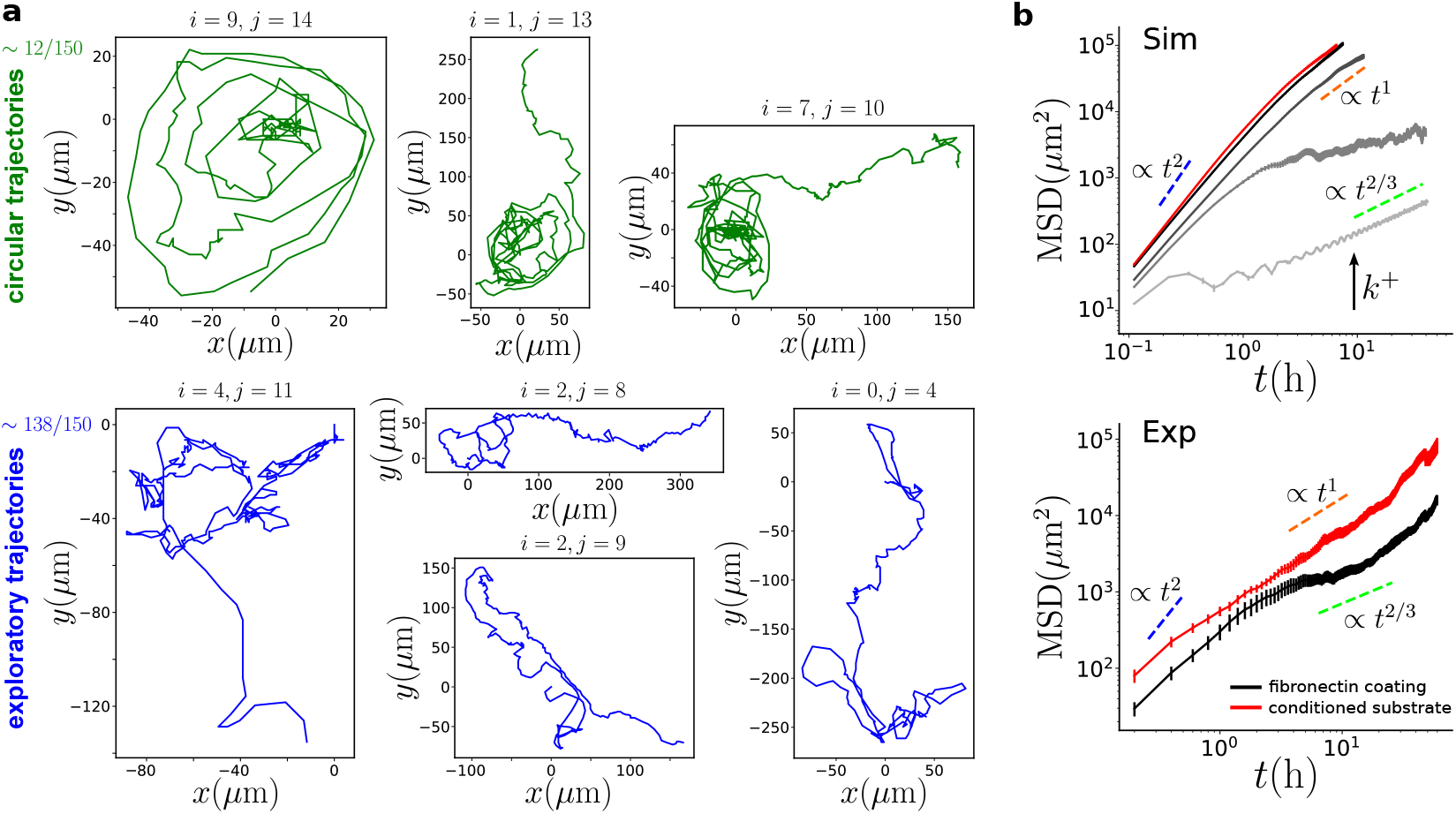
**a** Trajectories of MDCK cells plated on fibronectin-coated micropatterned squares from (15). At the top, are examples of circular trajectories with increasing radii. In the cases shown in the middle and right panels, the cells eventually leave their circular paths and become more exploratory. At the bottom, are examples of persistent trajectories. Indices *i* and *j* are the coordinates for the trajectories in Fig. S3. Of 150 classified trajectories, 12 were circular and 138 were exploratory. **b** (Top) Mean Square Displacement (MSD) for *n* = 100 simulated trajectories for increasing values of *k*^+^ represented from lighter to darker grayscale, (*k*^+^ = { 0.1, 0.2, 0.3, 1.0} *c*_0_ */*s). The red curve corresponds to the conditioned substrate where the cells are placed on a surface saturated in footprint (*ψ*(**r**, *t*) = 1). (Bottom) Reproduction of the measured MSD for the position of cell *i* at time *t*, and *i* an index that runs over the *N* different trajectories. the MDCK cells from (15). The colored dashed lines represent different slopes in the log scale plot corresponding to sub-diffusion (slope=2/3, green), diffusion (slope=1, orange), and ballistic (slope=2, blue). We compute the MSD from an ensemble average of the squared displacements, 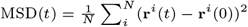, where **r**^*i*^(*t*) is the position of cell *i* at time *t*, and *i* an index that runs over the *N* different trajectories.

We quantify the degree of exploratory behavior by computing the mean square displacement (MSD) for both simulation and experiment in Fig. 7. In our simulations, below the transition, where cells move in circles with increasing radius, the MSD grows slowly as ∝ *t*^2*/*3^. This exponent corresponds to a cell that increases the explored area as 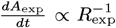, where *A*_exp_ is the explored area, *i*.*e*. the footprint area, and *R*_exp_ is the radius of the explored area assuming is a perfect circle. On the other end, for large values of *k*^+^, where cells move almost persistently until they reach the box edge, we observe a MSD that grows as ∝ *t*^2^ – cells are essentially ballistic. We also simulate a conditioned substrate (one that has had cells extensively crawling on it, and has saturated levels of footprint) by setting *ψ*(**r**, *t*) = 1 (red curve in Fig. 7**b** top panel), finding a similar ballistic dynamics. Towards the end of the simulated time range, the MSD for these high-deposition-rate cases starts to curve downwards to ∝*t*^1^ indicating a transition between a ballistic phase to a diffusive phase, which we don’t fully observe due to the size limitation of the simulation box. Interestingly, for the case of *k*^+^ slightly above the transition, which we think is the experimentally relevant case, the MSD shows an exponent less than 1, reflecting once again, that the trajectories spend a long time looping over the past traces, in the form of weak confinement. Though we do not see this due to the limited time of the simulations, we expect the subdiffusive behavior at *k*^+^ slightly above the transition will become diffusive at long enough times, as cells escape from loops and explore new regions.

The experimental data of (15) on MDCK cells on two-dimensional substrates shows an exponent below 1 at intermediate times, and a transition to more normal diffusion at longer times (Fig. 7**b** bottom panel). At intermediate times, this is consistent with what we observe at *k*^+^ = 0.2*c*_0_*/s* in our simulations (Fig. 7b). The experimentally-observed long-time change to diffusion could arise from cells escaping their loops, as we mentioned above. However, it could also potentially arise from cell-to-cell variability, as revealed in our classification of the experimental trajectories. For instance, imagine that the experiment consists of cells in two states with different parameters, leading to the MSD being a weighted average of the MSDs we see in simulation. If some cells (“exploratory”) have a long-time MSD∼*t*^1^ (e.g. due to high deposition rate) and others (“confined”) have a long-time MSD ∼*t*^*α*^ with *α <* 1, we would expect at long enough times MSD∼ *t*^1^ but at intermediate times could get something closer to *t*^*α*^. However, to get the intermediate *t*^*α*^ seen in experiments requires exploratory cells to have a lower MSD than confined cells at short times, so that the confined cell MSD is dominant. This could arise if cells with higher deposition rates also have slower motilities – trading off speed and ability to explore new territory. We calculated the MSD excluding circular trajectories, but did not observe a significant change in the shape of the MSD. This does not, however, rule out cell-to-cell variability as relevant, because our screening only catches those cells that make circular motions and not those that are subdiffusive in other ways as *e*.*g. k*^+^ = 0.2*c*_0_*/s*.

The combination of the observation of oscillation in 1D, the mixture of expanding circular and exploratory motion, and the MSD showing a transient subdiffusion are all consistent with our hypothesis that the MDCK cells observed in (15) are slightly above the transition to exploratory motion in 2D but below the transition in 1D.

## Discussion

In our simulations, we find sharp transitions between the different cell behaviors like confinement, persistent motion, and oscillations as a function of parameters like footprint deposition rate *k*^+^, sensitivity to footprint *K*_*D*_, and cell shape. This suggests that cells could achieve dramatic changes in their behavior by small regulations of their sensing machinery (32, 33, 41) or mechanics. In addition, if there is cell-to-cell variability in, e.g. deposition rate or response to the haptotactic cue *K*_*D*_, then different cells in an isogenic population could have vastly different behaviors. Consistent with this idea, in our re-analysis of the experiments of (15), we found evidence that cells on 2D substrates can develop both the confined, expanding circular behavior we predict at low deposition rate and also a more exploratory migration. We argue that MDCK cells in the experiments of (15) are tuned very close to the transition point between confined and exploratory behavior, being in most cases, slightly above it, with cell-to-cell variation leading to a few examples of confined migration. MDCK cells may benefit from being able to drastically vary their behavior with small regulations in their mechano/biochemical machinery and this – speculatively – suggests that they may have evolved to be at this optimal point.

Beyond the good agreement with the findings in (15), our model can be adapted to diverse cell types and ECMs. Distinctive elements such as cell mechanics, biochemistry, footprint deposition and response, as well as the interaction with the microenvironment, can be captured by adjusting the model parameters. Migration modes depend on cell type and ECM (8). For instance, less integrin-dependent migration modes may not respond as strongly to a footprint, leading to a lower *k*_*b*_ or higher *K*_*D*_. On the other hand, stiffer ECMs increase the strength of focal adhesions and cell mechanotransductive response, leading to higher traction forces and increased Rac1 activity– e.g. a higher protrusion coefficient, *α*, and Rac1 activation rate, *k*_*b*_. These quantitative changes may also explain the presence of oscillations in response to perturbations. For instance, in fibrosarcoma cells, depletion of zyxin induces a transition in cell motility behavior from random motion to oscillations in 1D and 3D media (38). Zyxin is a protein associated with focal adhesion sites playing a role in mechanotransduction. In terms of our model, we might think of zyxin depletion or other perturbations to focal adhesions as affecting how cells respond to the footprint. This could alter the parameter *K*_*D*_. When zyxin is depleted, the cell may respond less to the footprint, resulting in a larger *K*_*D*_ value. This increase in *K*_*D*_ could lead a cell to transition from persistent to oscillatory motion, as shown in Fig 3**c**.

Our results also explain why, even though footprints can have immediate and obvious effects on cells on micropatterned stripes (15), this effect was only discovered relatively recently. Cells that develop oscillations on microstripes may have quite simple behavior on 2D substrates, with a MSD showing the stereotyped transition from ballistic to diffusive (42) (Fig. 6-7, *k*^+^ = 0.3*c*_0_*/s*). We argue that many cell types may be more strongly affected by footprint deposition than currently thought, but that footprint effects are not apparent on 2D substrates or at high deposition rates. As a result, identifying the presence of footprint interactions from experimental cell trajectories can be challenging, leading to the underreporting of this phenomenon. This is consistent with the recent broader understanding of the relevance of the cell’s environment to its migration (8, 43–45). Footprint interactions could be tested by systematically studying different cell types on 1D and 2D substrates to determine whether other cell types exhibit similar motility behaviors, *i*.*e*., consistent with being slightly above the transition to exploratory movement, as we have found here.

We have a number of immediate predictions from our model. We identified the key factors of transitioning between oscillatory and persistent behavior as the rate of deposition of footprint and the threshold for responding to the footprint (Fig. 3**c**). Knockdowns of integrins specific to the footprint would then, we expect, make the cell less responsive to the footprint, akin to reducing *K*_*D*_ – leading to more persistent motion. Knockdown of other focal adhesion proteins might also play a role – as seen in the oscillatory behavior induced by zyxin knockdown (38). Haptotaxis may also be suppressed by knockdown of Arp2/3 (46) – though this is a dramatic intervention. In addition, we predict that degradation of the footprint over time will lead to cell oscillations reaching a fixed size, with the amplitude dependent on the degradation rate (Fig. 4**b**, Eq. (9)). While the footprint does not naturally degrade over a relevant timescale in (15), it might be possible to *induce* degradation controllably by introducing MMPs into the experiment. This would allow direct control over the steady-state amplitude of oscillation. We would also predict that in the presence of this degradation, cells would reach the same amplitude of oscillation independent of whether they are seeded on a normal fibronectin substrate or one that has been “conditioned” by having other cells crawl on it (Fig. 4b). Unfortunately, tests of these predictions require a clear identification of the essential components of the footprint – something which remains in progress.

In our model, we assumed that the footprint increases positive feedback of Rho GTPase at the front of the cell, such as Rac1 or Cdc42. We expect alternate models, e.g., that the footprint down-regulates de-activation of Rho GTPase *k*_*c*_ through inhibition of Rho GTPase GAPs, to give similar cell-footprint interactions. However, different feedback modes between footprint and the Rho GTPase might make quantitative changes in the threshold where cells respond to the end of the footprint. Quantitative changes in sensitivity to repolarization have been found for a related question – how cells respond to optogenetic stimulation affecting different parts of the Rho GTPase system (47). We have also neglected a possible feedback where changes in cell polarity alters deposition(34), instead keeping the deposition rate constant for simplicity.

In (15), oscillations were observed in MDCK, Caco-2, and RPE1 epithelial cells, although findings were less conclusive in the latter case. These are all standard epithelial cell lines, and the context of epithelial cells on fibronectin may be relevant to wound healing where fibronectin is a temporary matrix for the healing wound (48). MDCK is in particular a standard for measurements of single-cell, collective, and clustered motility (15, 49–52) and allows for straightforward measurement of Rac1 activity (15, 49). The presence of oscillations in cell types of different origin (immortalized canine kidney, colon carcinoma, healthy human retinal pigment cells), as well as later discoveries, suggest that this phenomenon is robust and generalizable. Others have also recently observed similar oscillatory behavior induced by modification of the surrounding ECM. Human bronchial epithelial cells plated under matrigel showed oscillatory motion as they slowly expanded their explored regions by interactively weakening and carving the ECM (53). Similarly, Th1 T cells deposited on polyacrylamide gels, when covered with agarose gel, migrated along a trajectory, expanding it with each pass as they moved back and forth (54). These findings suggest that the migration modes described here are not necessarily unique to our proposed mechanism or the particular experimental setup of (15) but may arise more generally from how cells make their way through modifying and responding to their environment.

We are not aware of the presence of single-cell oscillation or expanding circular trajectories being observed *in vivo*, but there is clearly evidence that deposited footprints are relevant *in vivo*, in particular in trails created by neutrophils (9). Oscillations have also been observed in more-realistic three-dimensional extracellular matrices (38, 53). As we have seen clear evidence in our modeling that the types of migration cells undergo are influenced both by the footprint properties and the cell’s geometric confinement, we would expect a wide spectrum of motility patterns *in vivo*, depending on the structure of the cell’s environment. Earlier work has shown complex migration *in vivo* including anisotropy, transitions between persistent and random motion, and broad distributions of cell step sizes (6, 55, 56).

We have assumed that the concentration of footprint deterministically drives Rac1 activation (Eq. (6)). However, at low concentrations of footprint ligand, the ability of cells to follow deposited signals may be impaired by physical sensing limits arising from stochasticity of ligand-receptor interactions, as extensively studied for chemotaxis, concentration sensing, and other relevant sensing problems (57–63), including haptotaxis (64). We suspect this noise is unlikely to be relevant in the results presented here – given the long timescale of cells making decisions and the sharp transitions in footprint concentration, we expect the relevant signal-to-noise ratio to be quite large. However, these concerns may be more relevant for cells attempting to distinguish old footprints left by previous cells, where degradation will reduce the signal.

The interplay between cells and footprint may be relevant in collective cell migration, where leader cells can pave the way for other cells to follow in their traces. This idea has recently been validated in *in vitro* experiments in which it has been observed that immune Th1 T cells show a preference for following the tracks that other T cells leave on the ECM (54). In cancer metastasis, a first line of cancer cells migrating away from a tumor could facilitate the way for others (65–68). Spatial footprints might also provide an alternative mechanism for cells to guide themselves through complex environments, analogous with work on self-generated gradients (14, 69). Our results suggest that these collective behaviors might also have many possible motility behaviors, which are under control of the factors we identify here.

### Parameters

## ACKNOWLEDGMENTS

EPI and BAC are supported by NIH R35 GM142847. This work was carried out at the Advanced Research Computing at Hopkins (ARCH) core facility (rockfish.jhu.edu), which is supported by the National Science Foundation (NSF) grant number OAC 1920103. This work was supported by LABEX Who Am I? (ANR-11-LABX-0071 to BL and JDA), the Ligue Contre le Cancer (Equipe labellisée 2019 to BL), the Agence Nationale de la Recherche (‘POLCAM’ ANR-17-CE13-0013, “Myofuse” ANR-19-CE13-0016), INCa 2018-1-PL BIO-08-ICR-1 (Decision N° 2018-154) and DIM Elicit “Région Ile-de-France”. We acknowledge the ImagoSeine core facility of the IJM, member of IBiSA and France-BioImaging (ANR-10-INBS-04) infrastructures. We thank Pedrom Zadeh and Wei Wang for a close reading of the draft.

