## Supplementary Information for "Deposited footprints let cells switch between confined, oscillatory, and exploratory migration"

Emiliano Perez Ipiña[*]

*Department of Physics & Astronomy,*

*Johns Hopkins University, Baltimore, MD.*

Joseph d'Alessandro[†] and Benoît Ladoux[‡]

*Université Paris Cité, CNRS, Institut Jacques Monod, F-75013 Paris, France*

Brian A. Camley[§]

*Department of Physics & Astronomy and Biophysics,*

*Johns Hopkins University, Baltimore, MD.*

[*]

[†]

[‡]

[§]

### S1. SUPPLEMENTARY FIGURES

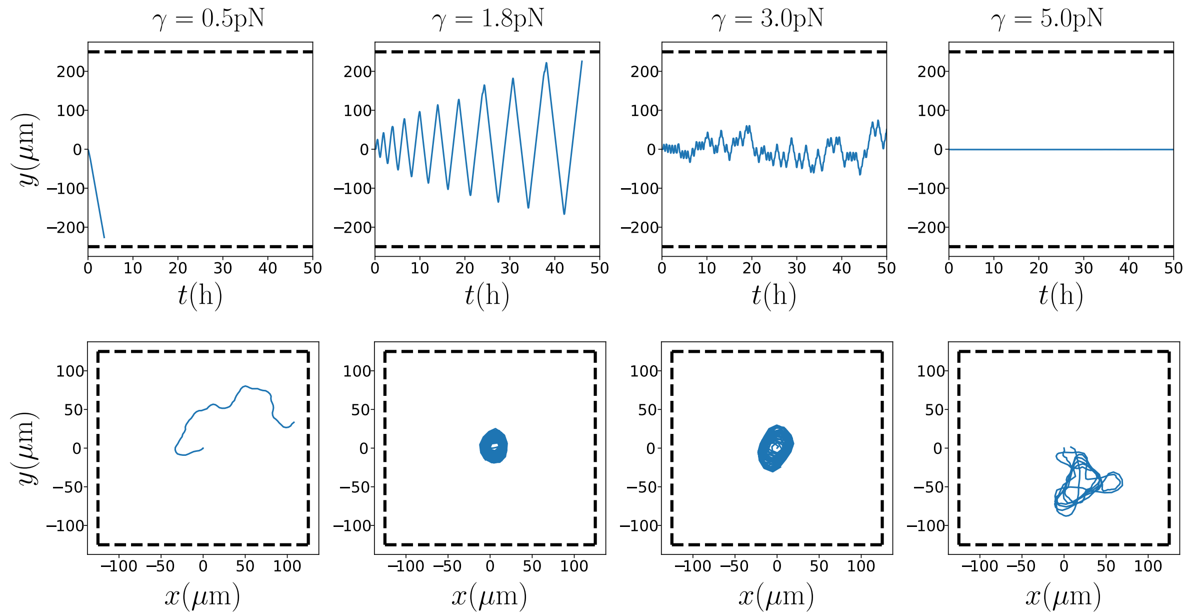


FIG. S1. Transitions between confined and persistent/exploratory motion depend on cell shape. Simulated cell trajectories for different cell membrane tension coefficient values $\gamma$ for 1D stripes (top) and 2D (bottom). The membrane tension coefficient determines how easily cells can be deformed and vary their shape. A low value of $\gamma$ indicates that cells can be easily deformed, while a high value of $\gamma$ means that the cell membrane is resistant to deformation, resulting in more round-shaped cells. The default value for the tension coefficient used along the results of this paper is $\gamma = 1.8\,$pN. We used $k^+ = 0.3\,c_0\text{s}^{-1}$ for 1D stripes and $k^+ = 0.1\,c_0\text{s}^{-1}$ for 2D substrates.

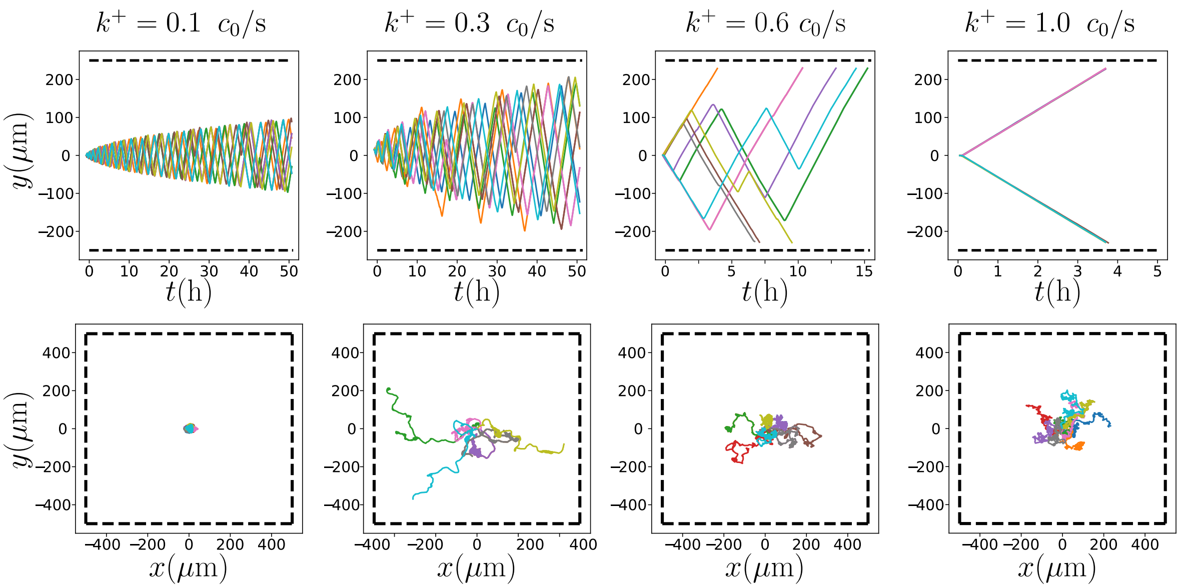


FIG. S2. Noise in cell polarity affects their trajectories. We show simulations of trajectories with the same conditions of Fig. 6 but with a higher polarity noise, $\sigma_\rho = 0.38\,\mu\mathrm{m}^{-2}\,\mathrm{s}^{-1/2}$. Trajectories in 1D stripes are robust to polarity noise. Cells occasionally reverse polarity before reaching the edge or can move forward a few microns further.

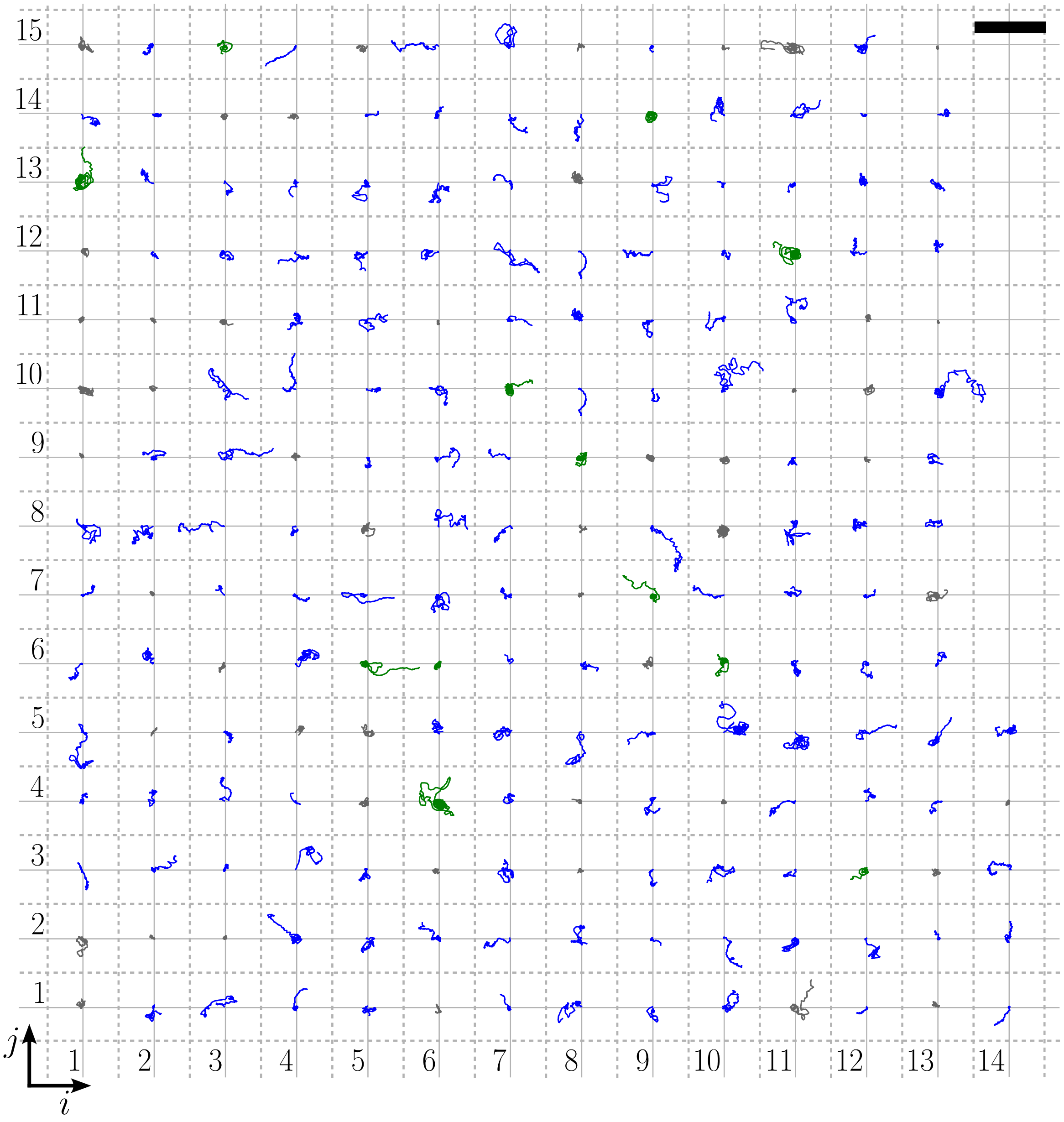


FIG. S3. Array of all the experimental trajectories of MDCK cells on the 2D squared substrates coated with fibronectin ($n = 200$). The trajectories were classified using an initial automated selection method followed by eye inspection into circular (12/150 – green), and exploratory (138/150 – blue). Of the 200 trajectories, there are 50 that we could not confidently identify to which category they belong. We represented them in gray. For more detail on the classification method see Section S3 in this SI. The scale bar represents 500$\mu$m.

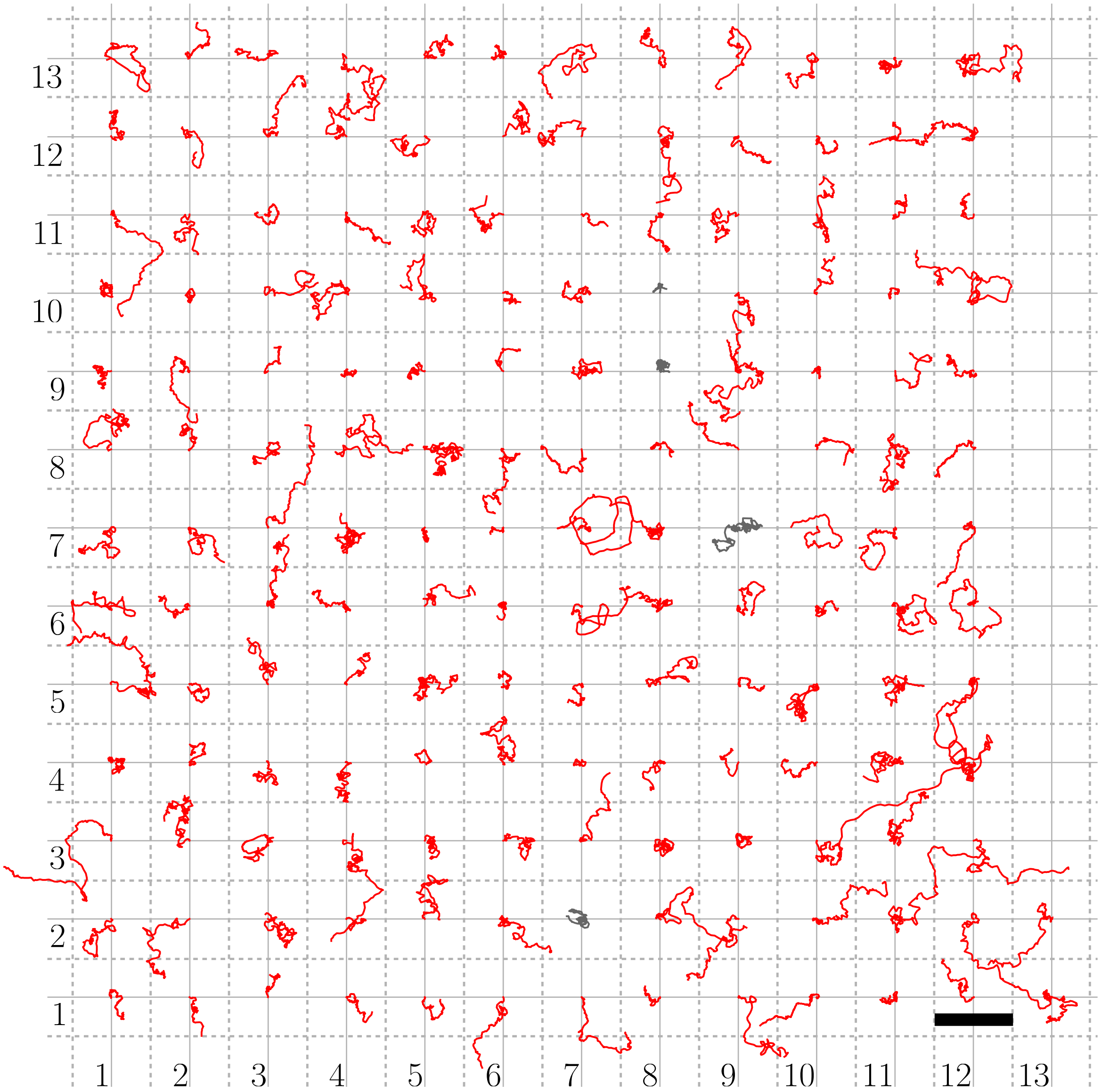


FIG. S4. Same as Fig. S3 but for cells placed over the conditioned substrate. A blind classification from a random mixture of conditioned and unconditioned substrate trajectories yielded 154 exploratory (red), 0 circular and 4 not classified (gray) conditioned substrate trajectories out of 158. The scale bar represents 500$\mu$m.

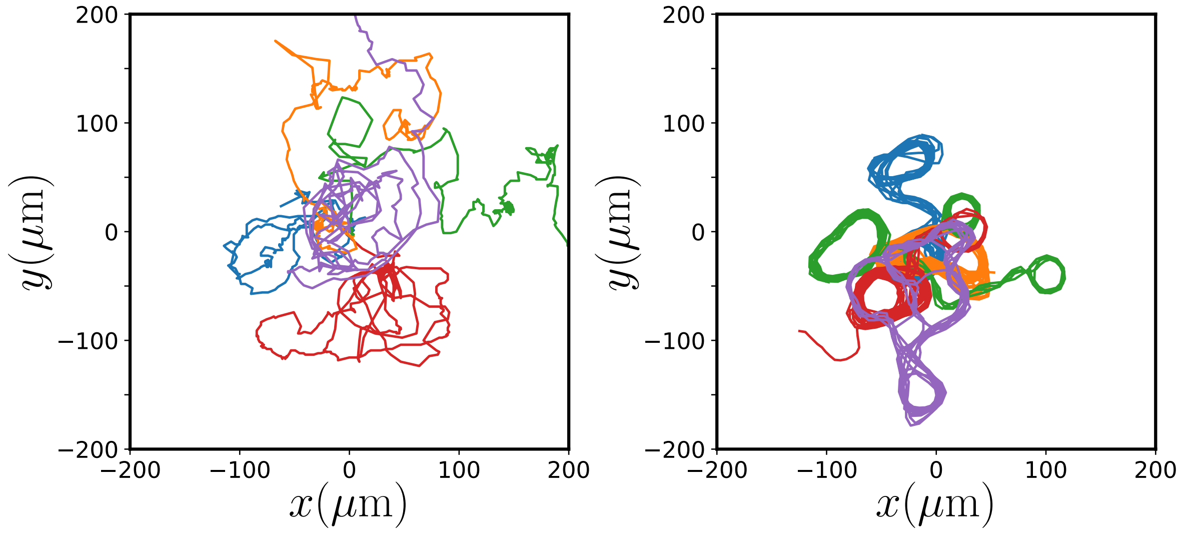


FIG. S5. Examples of 5 trajectories for both experiments and simulations. On the left, trajectories of MDCK cells placed on a 2D square initially coated with fibronectin from [1]. On the right, simulated trajectories of cells placed under the same conditions as in the experiment. Deposition rate $k^{+} = 0.2\, c_0 \mathrm{s}^{-1}$. Experimental trajectories show that cells exhibit a combination of motility behaviors in which they spend some time exploring new areas while tending to remain close to previous trajectories. This is similar to what is observed in the simulated trajectories – though the experimental trajectories are noisier.

### S2. NUMERICAL IMPLEMENTATION OF THE MODEL

To solve our mathematical model we integrate the evolution equations of $\phi(\mathbf{r},t)$, $\rho(\mathbf{r},t)$ and $c(\mathbf{r},t)$ numerically. We use a uniform grid ($x-$axis $\times$ $y-$axis) of 50 $\times$ 1000 points for the simulation over the stripe micropatterns (1D) and 500 $\times$ 500 (Fig. 5) and 2000 $\times$ 2000 (Figs. 6-7)points in the case of the square micropatterns (2D). The spatial grid size was set the same for both axes, $\Delta r = \Delta x = \Delta y = 0.5\mu$m. Then, we implement the notation $\mathbf{r} = (x_i - x_c, y_j - y_c)$, where $i$ and $j$ are indexes representing the lattice site coordinates and $(x_c, y_c)$ in the position of the grid center.

We implemented an explicit finite difference numerical method to solve the phase-field evolution, $\phi(\mathbf{r},t)$. From Eqs. (1)-(2) on the main text we need to solve

$$\partial_t\phi(\mathbf{r},t) = \frac{1}{\tau}\left(\alpha\rho(\mathbf{r},t) - \beta\right)|\nabla\phi| + \frac{\gamma}{\tau}\left(\nabla^2\phi - \frac{G'(\phi)}{\epsilon^2}\right), \tag{S1}$$

where $G'(\phi) = \frac{\mathrm{d}G}{\mathrm{d}\phi} = 36\phi\left(2\phi^2 - 3\phi + 1\right)$. For the time integration, we use a simple Euler forward method, $\partial_t\phi(\mathbf{r},t) \approx \frac{\phi(\mathbf{r},t+\Delta t)-\phi(\mathbf{r},t)}{\Delta t}$ with integration time step $\Delta t = 0.005$s. To compute the gradient modulus and the first-order spatial derivative we use a 5-point stencil,

$$\begin{aligned}
|\nabla\phi|(x_i,y_j) \approx& \sqrt{\Delta\phi_x^2(x_i,y_j) + \Delta\phi_y^2(x_i,y_j)},\\
\Delta\phi_x(x_i,y_j) \approx& \frac{\phi(x_{i-2},y_j) - 8\phi(x_{i-1},y_j) + 8\phi(x_{i+1},y_j) - \phi(x_{i+2},y_j)}{12\Delta r},\\
\Delta\phi_y(x_i,y_j) \approx& \frac{\phi(x_i,y_{j-2}) - 8\phi(x_i,y_{j-1}) + 8\phi(x_i,y_{j+1}) - \phi(x_i,y_{j+2})}{12\Delta r}.
\end{aligned}$$

where $x_{i-n} = x_i - n\Delta r$ and $y_{j-m} = y_j - m\Delta r$. Then, for the case of the Laplacian operator we used a 9-point stencil,

$$\begin{aligned}
\nabla^2\phi(x_i,y_j) \approx& \Big(\phi(x_{i+1},y_{j+1}) + \phi(x_{i+1},y_{j-1}) + \phi(x_{i-1},y_{j+1}) + \phi(x_{i-1},y_{j-1})\\
&+ 4\phi(x_{i+1},y_j) + 4\phi(x_i,y_{j-1}) + 4\phi(x_{i-1},y_j) + 4\phi(x_i,y_{j+1}) - 20\phi(x_i,y_j)\Big)\Big/6\Delta r^2.
\end{aligned}$$

In this way, the residual error for the spatial derivatives is $\mathcal{O}(\Delta r^4)$. Next, we can integrate all the terms in Eq. S1 and compute numerically the evolution of $\phi(\mathbf{r},t)$,

$$\begin{aligned}
\mathcal{T}_1(\mathbf{r},t) =& \frac{1}{\tau}(\alpha\rho(\mathbf{r},t) - \beta)|\nabla\phi|(\mathbf{r},t),\\
\mathcal{T}_2(\mathbf{r},t) =& -\frac{\gamma}{\tau\epsilon^2}\,G'(\mathbf{r},t),\\
\mathcal{T}_3(\mathbf{r},t) =& \frac{\gamma}{\tau}\,\nabla^2\phi(\mathbf{r},t),\\
\phi(\mathbf{r},t+\Delta t) =& \phi(\mathbf{r},t) + \Delta t\big(\mathcal{T}_1(\mathbf{r},t) + \mathcal{T}_2(\mathbf{r},t) + \mathcal{T}_3(\mathbf{r},t)\big).
\end{aligned}$$

We then solve the evolution of the active Rac1, $\rho(\mathbf{r},t)$, given by Eq. (5). We follow the same approach used in [2]. Applying the Euler-Maruyama forward method for time integration, we can write Eq. (5) in its discretized form,

$$\begin{aligned}\phi(\mathbf{r},t)\frac{\rho(\mathbf{r},t+\Delta t)-\rho(\mathbf{r},t)}{\Delta t}+\rho(\mathbf{r},t)\frac{\phi(\mathbf{r},t+\Delta t)-\phi(\mathbf{r},t)}{\Delta t} =&\nabla\cdot(\phi(\mathbf{r},t)D_\rho\nabla\rho(\mathbf{r},t))\\&+\phi(\mathbf{r},t)f_\rho(\mathbf{r},t)+\frac{\sigma_\rho\Xi(\mathbf{r},t)}{\Delta t}\end{aligned}$$

where $\Xi(\mathbf{r},t)=\int_t^{t+\Delta t}dt'\xi(\mathbf{r},t)$ is a Gaussian random number with zero mean and variance $\langle\Xi(\mathbf{r},\Delta t)\Xi(\mathbf{r}',\Delta t)\rangle=\Delta t\delta(\mathbf{r}-\mathbf{r}')$. From there we have,

$$\rho(\mathbf{r},t+\Delta t)=\frac{2\phi(\mathbf{r},t)-\phi(\mathbf{r},t+\Delta t)}{\phi(\mathbf{r},t)}\rho(\mathbf{r},t)+\Delta t\frac{\nabla\cdot(\phi(\mathbf{r},t)D_\rho\nabla\rho(\mathbf{r},t))}{\phi(\mathbf{r},t)}+\Delta t f_\rho(\mathbf{r},t)+\sigma_\rho\Xi(\mathbf{r},t), \tag{S2}$$

Note that Eq. (S2) has a dependence on both $\phi(\mathbf{r},t)$ and $\phi(\mathbf{r},t+\Delta t)$. The second term on the right of Eq. (S2) can be worked out to,

$$\mathcal{T}_2(\mathbf{r},t)=\Delta tD_\rho\frac{\nabla\phi(\mathbf{r},t)\cdot\nabla\rho(\mathbf{r},t)+\phi(\mathbf{r},t)\nabla^2\rho(\mathbf{r},t)}{\phi(\mathbf{r},t)}.$$

In the same manner, as we did for the integration of $\phi(\mathbf{r},t)$, here we also use a 5-point stencil for the first-order spatial derivatives of $\phi(\mathbf{r},t)$ and $\rho(\mathbf{r},t)$, and a 9-point stencil for the Laplacian of $\rho(\mathbf{r},t)$. We then can break down Eq. (S2) in the following four terms,

$$\begin{aligned}\mathcal{T}_1(\mathbf{r},t)=&\frac{2\phi(\mathbf{r},t)-\phi(\mathbf{r},t+\Delta t)}{\phi(\mathbf{r},t)}\rho(\mathbf{r},t),\\\mathcal{T}_2(\mathbf{r},t)=&\Delta tD_\rho\left(\Delta\phi_x(\mathbf{r},t)\Delta\rho_x(\mathbf{r},t)+\Delta\phi_y(\mathbf{r},t)\Delta\rho_y(\mathbf{r},t)+\phi(\mathbf{r},t)\nabla^2\rho(\mathbf{r},t)\right)\Big/\phi(\mathbf{r},t),\\\mathcal{T}_3(\mathbf{r},t)=&\Delta tf_\rho(\mathbf{r},t),\\\mathcal{T}_4(\mathbf{r},t)=&\sigma_\rho\Xi(\mathbf{r},t).\end{aligned}$$

Then we compute these four terms to finally obtain the evolution of $\rho(\mathbf{r},t)$,

$$\rho(\mathbf{r},t+\Delta t)=\mathcal{T}_1(\mathbf{r},t)+\mathcal{T}_2(\mathbf{r},t)+\mathcal{T}_3(\mathbf{r},t)+\mathcal{T}_4(\mathbf{r},t).$$

Lastly, we must integrate Eq. (3). Here again, we use the Euler forward method for the time integration,

$$c(\mathbf{r},t+\Delta t)=c(\mathbf{r},t)(1-k^-\Delta t)+k^+\phi(\mathbf{r},t)\Delta t. \tag{S3}$$

As a note on the numerical implementation of the methods described above, to speed up the simulations, we only integrate the Eqs. (S1)-(S2) within a box of size $5R\times5R$ around the center of mass of the cell. We are allowed to do this since $\phi$ and $\rho$ rapidly drop to zero outside the cell's boundary.

We used a time step of $\Delta t = 0.05$s for 1D stripes and $\Delta t = 0.2$s for 2D substrates simulations. We performed verification simulations with $\Delta t = 0.002$s to check for numerical artifacts. We especially ensured the transition between confinement and exploratory motion was independent of $dt$ values. Simulation data were stored with a frequency $\Delta t_{\mathrm{data}} = 250$s and $\Delta t_{\mathrm{data}} = 400$s for the 1D stripes and 2D substrates respectively.

### S3. SELECTION AND CLASSIFICATION OF MDCK CELL TRAJECTORIES

We analyzed the experimental trajectories of MDCK cells placed on 2D square surfaces coated with fibronectin (US – unconditioned substrates), as well as on conditioned substrates (CS) from [1]. This is the original data from [1]'s Figure 4h. From that original sample, MDCK cell trajectories longer than 30h (+150 data points – data recorded frequency $\Delta t = 12$min) were considered yielding a total of 200 and 158 trajectories for US and CS, respectively.

Cell trajectories were classified according to their motility behavior into circular or exploratory. The classification process was done in two stages. We first selected those trajectories that were candidates for circular motion. These trajectories are characterized by either not moving too far from their origin **or** by having a large traveled path compared to their distance from their origin – i.e. those that moved a lot without having a large net displacement, as would be expected for circular motion. For the first condition, we selected those trajectories where the maximum distance to their origin, $r_{\max} = \max(\sqrt{(x-x_0)^2 + (y-y_0)^2})$, was less than $50\mu$m. For the second condition, we computed the traveled path, $l(t_k) = \sum_{i=1}^{k-1} \sqrt{(x(t_{i+1}) - x(t_i))^2 + (y(t_{i+1}) - y(t_i))^2}$, where $i$ and $k$ are indexes that run over the data points. Then, we made a linear fit of $l(t_k)$ as a function of the distance from their origin, $r(t_k)$, and selected those trajectories with a slope greater than 4.5. This allows us to identify those trajectories that travel a significant distance without moving far from the origin. Trajectories that met one of the two conditions were selected as candidates for circular motion and passed to the second stage. Trajectories not selected were classified as exploratory. In the second stage, we manually examined the selected trajectories and sorted them into their respective categories by visual inspection. To this end, we considered simultaneously plots of $x - y$ with the raw data and a moving average of $n = 5$ points, as well as plots of $x$ and $y$ over time. In certain cases, the trajectories showed some ambiguity as to the category to which they belonged: we could not say with certainty whether they were circular or exploratory. In other cases, the cells did not seem to be motile. These trajectories were left out of the classification and labeled as "not classified".

To verify that the circular behavior is specific to cells moving in the unconditioned substrate and not a common motility pattern, we performed a blind classification of the trajectories. Thus, we mixed and shuffled the US and CS data so it was not possible to know in advance to which experiment each trajectory belonged. Then, we applied the classification method described above over the total amount of 358 trajectories. The initial algorithmic filter selected 100 candidates for circular motion. After blind classification of these candidates, we finally obtained 12 circular and 138 exploratory trajectories for the unconditioned substrate case, and 0 circular and 154 exploratory for the conditioned substrate one. Of the 100 candidate trajectories 54 were labeled as not classified, – 50 for the US and 4 for the CS.
